# pyActigraphy: open-source python package for actigraphy data visualisation and analysis

**DOI:** 10.1101/2020.12.03.400226

**Authors:** Grégory Hammad, Mathilde Reyt, Nikita Beliy, Marion Baillet, Michele Deantoni, Alexia Lesoinne, Vincenzo Muto, Christina Schmidt

## Abstract

Over the past 40 years, actigraphy has been used to study rest-activity patterns in circadian rhythm and sleep research. Furthermore, considering its simplicity of use, there is a growing interest in the analysis of large population-based samples, using actigraphy. Here, we introduce *pyActigraphy*, a comprehensive toolbox for data visualization and analysis including multiple sleep detection algorithms and rest-activity rhythm variables. This open-source python package implements methods to read multiple data formats, quantify various properties of rest-activity rhythms, visualize sleep agendas, automatically detect rest periods and perform more advanced signal processing analyses. The development of this package aims to pave the way towards the establishment of a comprehensive open-source software suite, supported by a community of both developers and researchers, that would provide all the necessary tools for in-depth and large scale actigraphy data analyses.

**Required Metadata:** *Current code version:* 

## 1. Motivation and significance

Actigraphy consists in continuous movement recordings, using small watch-like accelerometers that are usually worn on the wrist or on the chest. As recordings can last several days or weeks, this technique is an adequate tool for in-situ assessments of the locomotor activity and the study of rhythmic rest-activity patterns. Consequently, it has been used in the field of sleep and circadian rhythm research [1] to assess night-to-night variability in estimated sleep parameters as well as rest-activity rhythm integrity. For example, intradaily variability has been associated with both cognitive and brain ageing [2, 3], while sleep fragmentation, as quantified by probability transitions from rest to activity during night-time, has been linked to cognitive performances [4] as well as to increased risks for Alzheimer’s disease [5].

However, the generalization of the findings made by this technic remains difficult; researchers either develop specific, often closed-source, data processing pipeline and/or analysis scripts, which are time-consuming, error prone and make the reproducibility of the analyses difficult, or they rely on commercial toolboxes that are not only costly but also act as black boxes. In addition, cumbersome manual data preprocessing, such as cleaning, hampers large scale analyses, which are mandatory for reliable and generalizable results.

**Table 1:** Code metadata (mandatory)

| Nr. | Code metadata description | Please fill in this column |
| --- | --- | --- |
| C1 | Current code version | v1.0 |
| C2 | Permanent link to code/repository used for this code version | https://github.com/ghammad/pyActigraphy |
| C3 | Code Ocean compute capsule |  |
| C4 | Legal Code License | GPL-3.0 |
| C5 | Code versioning system used | Git |
| C6 | Software code languages, tools, and services used | Python 3 |
| C7 | Compilation requirements, operating environments & dependencies |  |
| C8 | If available Link to developer documentation/manual |  |
| C9 | Support email for questions |  |

In 2012, the UK Biobank decided to add 7-day actimetry-derived physical activity data collection. However, only a reduced set of sleep estimates has been extracted yet from this dataset to identify different rest-activity phenotypes and link them to pathology of genetic background (e.g. [6, 7]).

We thus argue that there is a need for a comprehensive and open-source toolbox for actigraphy data analysis. This motivated the development of the *pyActigraphy* package.

## 2. Software description and documentation

The *pyActigraphy* package is written in Python 3 (Python Software Foundation, https://www.python.org/) and is available from the Python Package Index (PyPI) repository. Its source code is hosted by Github and the Zenodo platform [8]. The online documentation of the *pyActigraphy* package contains a detailed description of the attributes and methods of its various modules and is meant to be used as complementary material of the current paper. In addition, more than a dozen of tutorials are made available online (https://ghammad.github.io/pyActigraphy/tutorials.html) to illustrate how to use the multiple features of the package, described in this paper. These tutorials are based on example data files that are provided with the package itself.

### 2.1. Reading native actigraphy files

The *pyActigraphy* package provides a unified way to read several actigraphy file formats. Currently, it supports files from:

- wGT3X-BT, Actigraph (.agd file format only);
- Actiwatch 4 and MotionWatch 8, CamNtech;
- ActTrust 2, Condor Instruments;
- Daqtometer, Daqtix;
- Actiwatch 2 and Actiwatch Spectrum Plus, Philips Respironics.

For each file format, a dedicated class has been implemented to extract the corresponding actigraphy data, as well as the associated meta-data. These classes inherit from a base class implementing the various functionalities of the *pyActigraphy* package. In addition, the package allows users to read actigraphy recordings, either individually, for visual inspection for example, or by batch, for analysis purposes.

### 2.2. Masking and cleaning data

Before analysing the data, spurious periods of inactivity, where the actigraph was most likely removed by the participant, need to be discarded from the activity recordings. The *pyActigraphy* package implements a method to automatically mask continuous periods of total inactivity. User-defined periods of masking can also be specified, either manually or in a specific file. In addition to temporary actigraph removals, another usual source of artificial inactivities arises when the recordings start before and/or end after the actigraph is actually worn by the participant. Upon reading an actigraphy file, the *pyActigraphy* package allows users to discard such inactivity periods by specifying a start and a stop timestamp. The data collected outside this time range are not analysed. These timestamps can also be specified by batch by using a simple log file where each line should correspond to the participant’s identification. This file is then processed to automatically apply such boundaries to the corresponding actigraphy file read by the package.

### 2.3. Activity profile and onset/offset times

In circadian rhythm and sleep research, profile plots of the mean daily activity of actigraphy recording provides a visual tool to assess the overall rest-activity pattern, as well as recurrent behaviours such as naps. This profile is obtained by averaging consecutive data points that are 24h apart, over the consecutive days contained in the recording. The *pyActigraphy* package provides methods to construct this profile (Fig 1). In addition, it provides two methods to calculate and anchor these 24-hour profiles to the average activity onset and offset times of a given individual in order to ease group averaging. These activity onset and offset times are defined as the time points where the relative difference between the mean activity before and after this time point is maximal and minimal, respectively.

**Figure 1:**
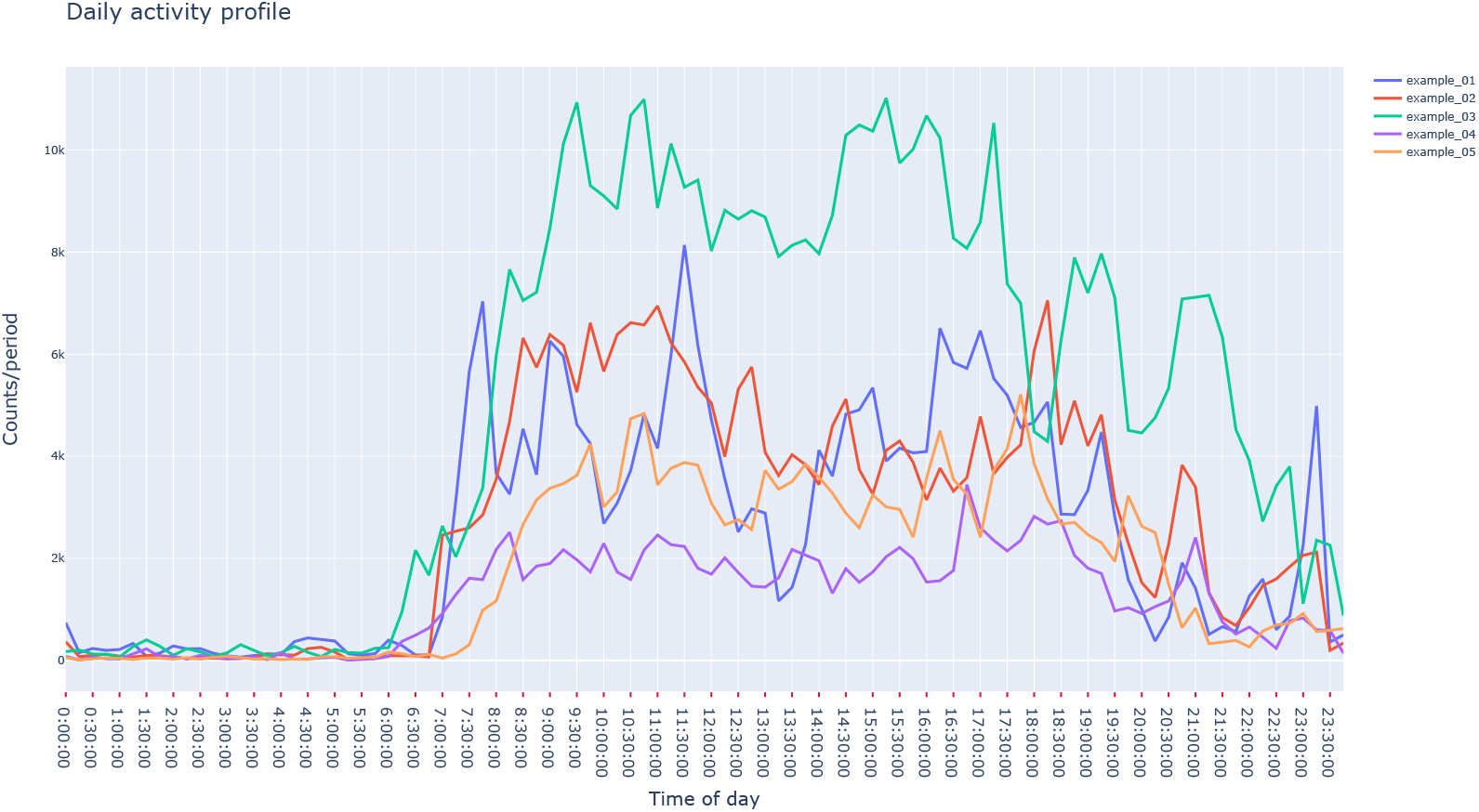
Vizualisation example of average daily profiles obtained with *pyActigraphy* using example files included in the package.

### 2.4. Visualization of sleep agenda

In both sleep research and medicine, a sleep diary is usually given with an actimeter to allow participants to report sleep episodes (duration and timing) as well as the subjective assessment of sleep quality for example. It allows comparisons between data recorded by an actigraph and the subjective perception of the individual wearing the device. In medical fields, sleep diaries are commonly recommended in order to help doctors in the diagnosis and treatment of sleep-wake disorders. The *pyActigraphy* package allows users to visualize and analyse sleep diaries, encoded as .ods or .csv files. Each row of these files indicates a new event, characterized by a type, a start time and an end time. A summary function provides descriptive statistics (mean, std, quantiles, …) for each type of events. For convenience and considering the current interests of the researchers involved in the development of the package, four types (active, nap, night, no-wear) are implemented by default when a sleep diary is read. However, the *pyActigraphy* package allows users to remove or customize these types and add new ones. As shown in Fig. 2, the visualization of the sleep diary is allowed through the use of the python plotting library “plotly” [9]. Each event found in the sleep diary is associated with a plotly “shape” object that can be overlaid with the actigraphy data in order to visually assess the adequacy between the subjective reports and their objective counterparts.

**Figure 2:**
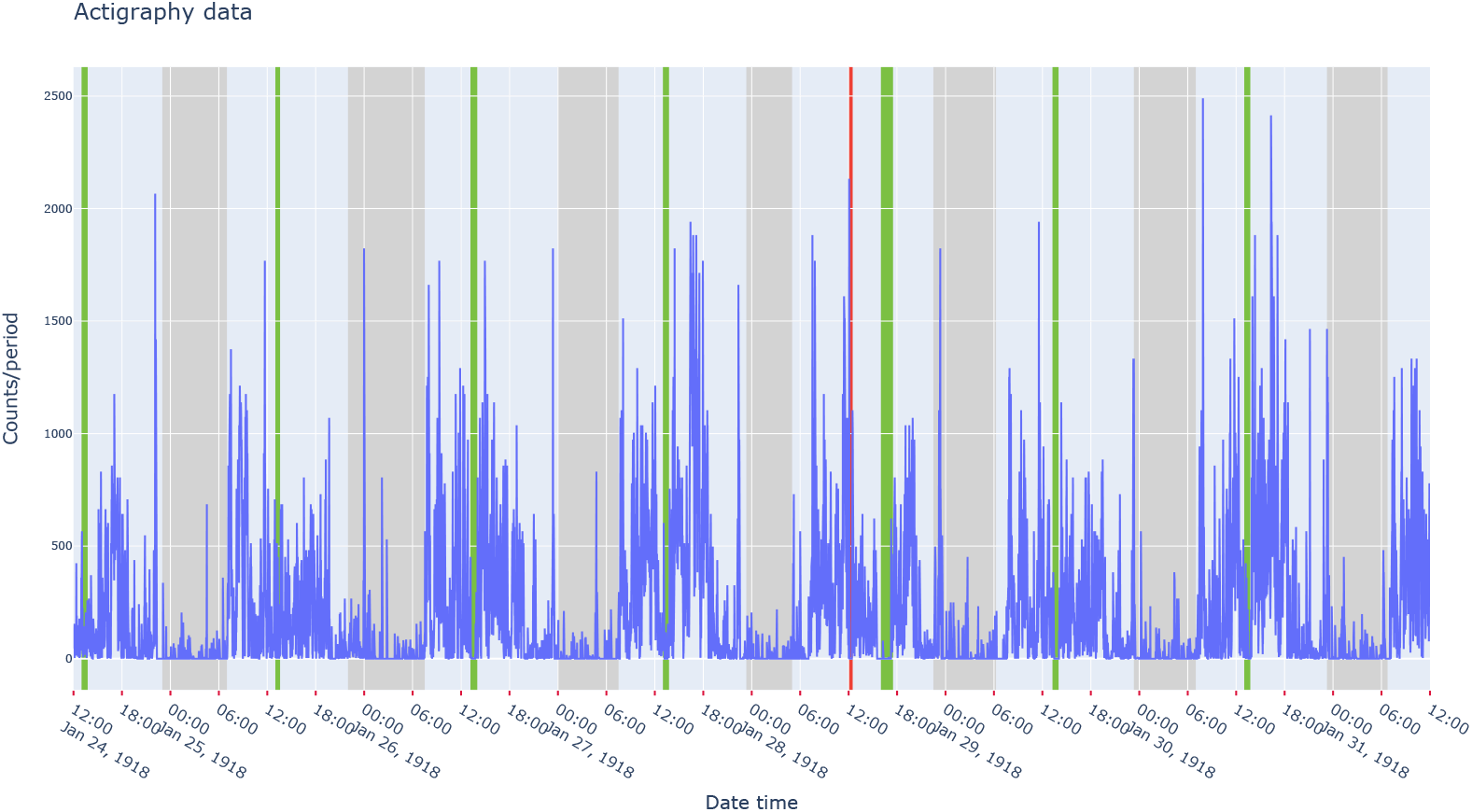
Vizualisation example of actigraphy data, overlaid with periods (green: nap, grey: night, red: device not worn) reported in the sleep diary example file included in the package.

### 2.5. Rest-activity rhythm variables

Non-parametric rest-activity variables can easily be calculated with the *pyActigraphy* package. The list of such variables includes:

- the interdaily stability (IS) and the intradaily variability (IV) [10], which quantify the day-to-day variance and the activity fragmentation, respectively;
- the relative amplitude (RA) [11], which measures the relative difference between the mean activity during the 10 most active hours (M10) and the 5 least active ones (L5).

In addition, *pyActigraphy* implements the mean IS and IV variables, namely ISm and IVm [12], obtained by averaging IS or IV values calculated with data resampled at different frequencies. Finally, the *pyActigraphy* package allows users to calculate the values of the IS(m), IV(m) and RA variables for consecutive, non-overlapping time periods of user-defined lengths. Upon calling the corresponding function, users can specify the resampling frequency, if the data must be binarized before calculation, as well as the threshold used to binarize the data.

### 2.6. Fragmentation of rest-activity patterns

The *pyActigraphy* package implements rest-activity state transition probabilities, *k*_*RA*_ and *k*_*AR*_ [13]. These variables quantify the fragmentation of the rest-activity pattern fragmentation; based on a probabilistic state transition model, where epochs with no activity are associated to a “rest” state (R) and to an “active” state (A) otherwise, the *k*_*RA*_ variable is associated with the probability to transition from a sustained “rest” state to an “active” state and the *k*_*AR*_ variable is associated with the probability to transition from a sustained “active” state to a “rest” state. The *pyActigraphy* package allows users to restrict the computation of the *k*_*RA*_ and *k*_*AR*_ variables to specific period of the day. For example, to target sleep periods, users may specify the activity offset and onset times (see section 2.3), as derived from individual activity profiles, as time boundaries. In the case of the *k*_*RA*_ variable, this would provide a quantification of the sleep fragmentation, adapted to a subject’s specific rest periods.

### 2.7. Rest-activity period detection

The *pyActigraphy* package implements several rest-activity detection algorithms, which can be classified into two broad classes:

- Epoch-by-epoch rest/activity scoring algorithms: Cole-Kripke’s [14], Oakley’s [15], Sadeh’s [16] and Scripps’ [17] algorithms. The idea underlying these algorithms is to convolve the signal contained in a sliding window with a pre-defined kernel. Most algorithms use gaussian-like kernels. If the resulting value is higher than a certain threshold, then the epoch under consideration, usually the one located at the centre of the sliding window, is classified as active and as rest, otherwise. Finally, the window is shifted forward by one epoch and the classification procedure is repeated.
- Detection of consolidated periods of similar activity patterns: Crespo’s [18] and Roenneberg’s [19] algorithms. These two algorithms are fundamentally different from the epoch-by-epoch scoring algorithm as they intend to detect, at once, consolidated periods of rest. One advantage of this class of algorithms is that it provides a start and a stop time for each period classified as rest.

As illustrated in Fig. 3, these algorithms have been implemented to return a binary time series: 0 being rest or activity depending on the definition made in the original article describing the detection algorithm.

**Figure 3:**
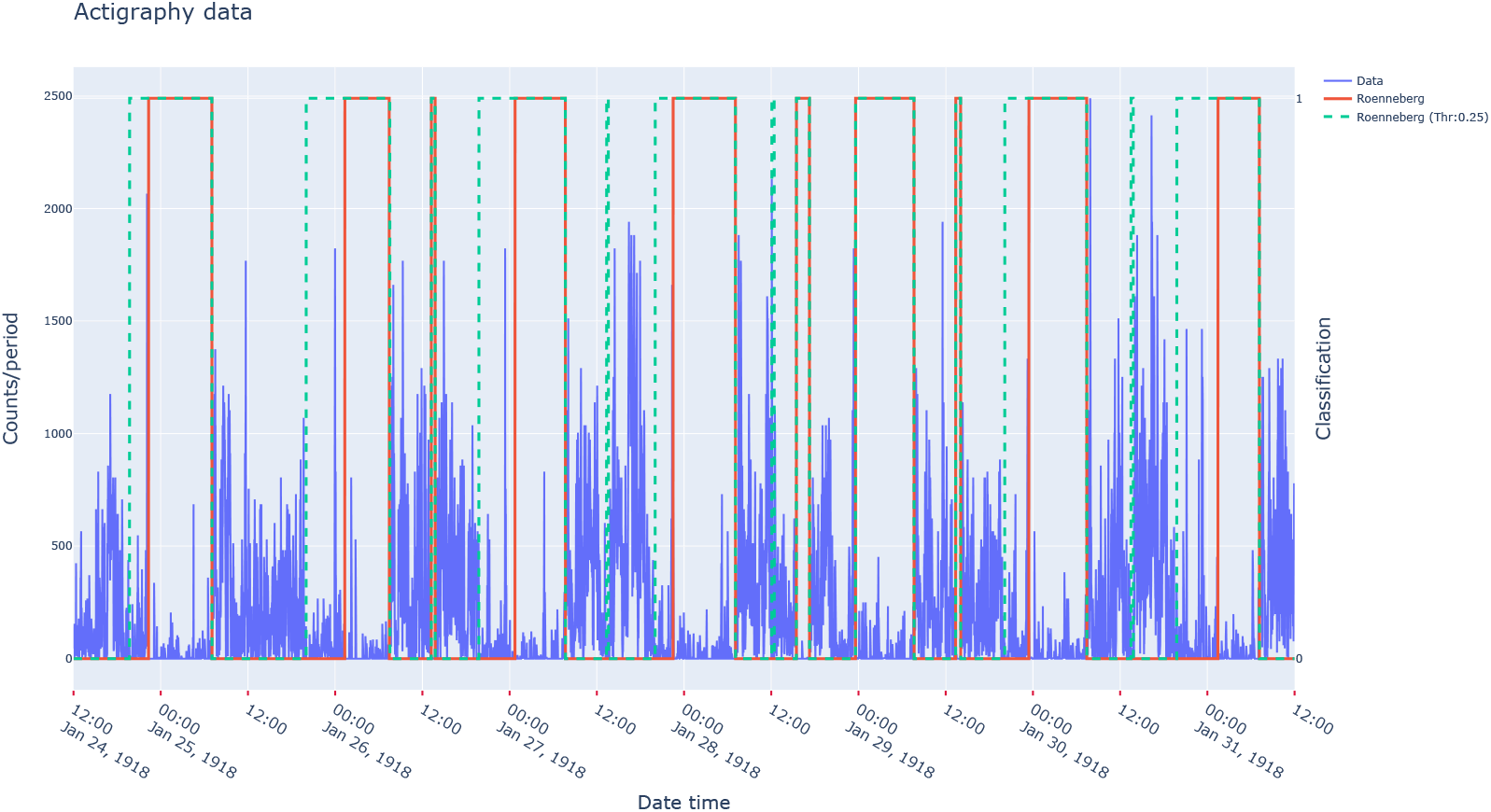
Vizualisation example of actigraphy data, overlaid with periods scored as “active” (0) or “rest” (1) by Roennberg’s algorithm [19] for two different settings (full line: default parameter values, dash line: with a threshold set at 0.25 of the activity trend).

Based on the aforementioned algorithms, the *pyActigraphy* package allows also the computation of a sleep regularity profile which quantifies the probability for the participant to be in the same state (rest or active) at any daytime point on a day-by-day basis. From this 24h profile, the sleep regularity index (SRI) [20, 21] can be calculated as the product of theses probabilities over all the time bins.

Finally, using the detection algorithms of the latter class, the *pyActigraphy* package allows the computation of the sleep midpoint as described in [21].

### 2.8. Advanced signal processing

The *pyActigraphy* package makes available additional functions for more advanced analyses of actigraphy recordings:

- Cosinor [22]: the idea of a Cosinor analysis is to estimate some key parameters of the actigraphy count series by fitting these data with a (co)sine curve:

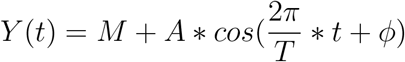

where M is the MESOR (Midline Statistic Of Rhythm), A is the amplitude of the oscillations, T is the period and *ϕ* is the acrophase. The fit procedure provides estimates of these parameters which can then help to characterize the 24h rest-activity rhythm of an individual.
- Detrented Fluctuation Analysis (DFA) [23, 24]: human activity exhibits a temporal organization characterised by scale-invariant (fractal) patterns over time scales ranging from minutes to 24 hours. This organization has been shown to be degraded with aging and dementia [25]. The DFA method allows the quantification of this scale-invariance and comprises four steps:
  1. Signal integration and mean subtraction
  2. Signal segmentation
  3. Local detrending of each segment
  4. Computation of the q-th order fluctuations
  All these steps have been implemented in the DFA class of *pyActigraphy*.
- Functional linear modelling (FLM) [26]: it consists in converting discrete measures to a function or a set of functions that can be used for further analysis. In most cases, the smoothness of the resulting function is under control, which ensures the derivability of this function. Three techniques are available in *pyActigraphy* to convert the actigraphy data to a functional form:
  – Fourier expansion
  – B-spline interpolation
  – Smoothing
  In the context of actigraphy, functional linear modelling and analysis have been successfully applied to link sleep apnea and obesity to specific circadian activity patterns [27].
- Locomotor inactivity during sleep (LIDS) [28]: the analysis of the locomotor activity during sleep revealed a rhythmicity that mimics the ultradian dynamic of sleep. This type of analysis opens new opportunities to study, *in situ*, sleep dynamics at a large scale and over large individual time periods. The LIDS class implements all the necessary functions to perform the analysis of the LIDS oscillations:
  – sleep bout filtering
  – non-linear conversion of activity to inactivity
  – extraction of the characteristic features of the LIDS oscillations via a cosine fit
- Singular spectrum analysis (SSA) [29, 30]: this technic allows the decomposition of a time series into additive components and the quantification of their respective partial variance. In the context of actigraphy, SSA can be used to extract the signal trend as well as circadian and ultradian components separately. The latter is relevant in human sleep research because sleep is not only alternating with wakefulness over the 24-hour cycle, but also exhibits an ultradian modulation, as mentioned previously. For example, a SSA analysis has been used to reveal alterations of the ultradian rhythms in insomnia [31]. All the necessary steps for the SSA and related functions, namely the embedding, the singular value decomposition, the eigentriple grouping and the diagonal averaging, are implemented in the SSA class. Since the subsequent calculations can be computationally intensive, the class implementation uses the open-source compiler Numba [32] for a direct translation of the functions to machine code and therefore improve their execution speed by several orders of magnitudes.

### 2.9. Online documentation and tutorials

The online documentation of the *pyActigraphy* package contains instructions to install the package, as well as informations about the authors and the code license. It also contains a detailed description of the attributes and methods available in the *pyActigraphy* package, which is generated automatically from source code annotations. In order to keep the documentation up to date with the latest developments of the package, the documentation is automatically generated anew and made available online for each new release. Finally, the online documentation offers several tutorials, illustrating the various functionalities of the package. These tutorials are generated from Jupyter notebooks [33] that are included in the *pyActigraphy* package itself, so that they can be used by any user to reproduce and practice the various functionalities of the *pyActigraphy* package in an interactive and user-friendly environment. As input data, the tutorials use real example data files that are included in the package for illustration and testing purposes. In total, 13 examples are included.

## 3. Illustrative Examples

As mentioned in section 2.9, the functionalities of the *pyActigraphy* package are illustrated in several notebooks that act as tutorials and are part of the online documentation. Nonetheless, this section provides two examples on how to read and analyse actigraphy files.

### 3.1. Basic example

The source code in Listing 1 is used to read multiple actigraphy files at once and calculate the rest-activity variables mentioned in sections 2.5 and 2.6. In this example, the results are simply printed but can be reused for further analyses.

Listing 1: Basic example

~~~
*#Import packages*
**import** pyActigraphy, os
*#Define path to example files*
*#(included in the pyActigraphy package)*
fpath = os. path.join(
        os. path.dirname(pyActigraphy. _ _file_ _), ‘tests/data/’
)
*#Read all Actiwatch 4 (CanNtech) files in the test directory:*
raw = pyActigraphy.io.read_raw(
        fpath+’example_0[0–9].AWD’,
        reader_type=’AWD’
)
*#Most functions can be accessed through this “raw” object.*
*#Ex: calculate non-parametric rest-activity variables*
ISs = raw.IS()
IVs = raw IV)
*#E: calculate the probability to transition*
*#from Rest to Active*
KRAs = raw.KRA(0)
~~~

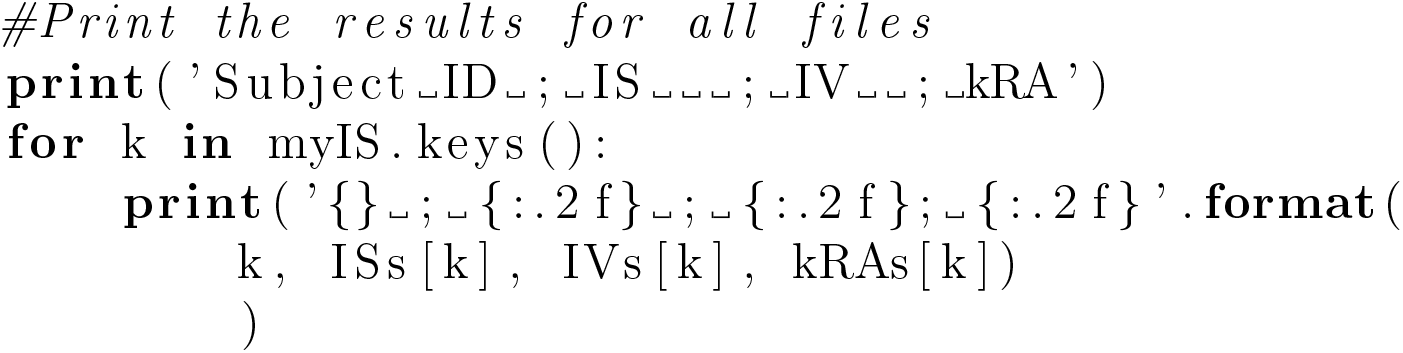

### 3.2. Advanced example

In Listing 2, a more complex example is provided. It illustrates how to fit actigraphy data with a cosinor model (section 2.8). In addition, the data are decomposed into several components via singular spectrum analysis (SSA) and the component whose pseudo-period is close to 24h is extracted.

Listing 2: Advanced example

~~~
*#Import packages*
**import** pyActigraphy, os
**from** pyActigraphy. analysis **import** Cosinor, SSA
*#Define path to an example file*
*#included in the pyActigraphy package)*
fpath = os. path.join(
        os. path.dirname(pyActigraphy. _ _file_ _), ‘tests /data/’
*#Read all Actiwatch 4 (ComNtech) files*
*#in the test directory:*
raw = pyActigraphy.io.read_raw_awd(fpath+’example_01 AWD’)
*#Initialize a Cosinor model object.*
my Cosinor = Cosinor()
*# and fit it to the data*
results = my Cosinor. fit (raw, verbose=True)
*#Inspect results*
results.params.pretty_print()
*# Cosinor model with parameters set to their estimated values*
cosinor_fit = cosinor. best_fit (raw, results .params)
*#Initialize a SSA object with the ‘raw’ object*
mySSA = SSA(raw. data, window_length=’24h’)
*# and fit it to the data*
mySSA. fit()
*#Inspect the singular values*
mySSA lambda_s
*#Calculate the weighted correlation matrix*
*#for the first 10 components*
w_corr_mat = mySSA. w_correlation_matrix (10)
*#Based on the results of the weighted correlation matria,*
*#it is straightforward to realize that*
*#the first and second SSA components*
*# (X_tilde) are strongly correlated and need to be merged.*
circ = mySSA. X_tilde ([1,2])
~~~

The result of the Cosinor model, as well the circadian SSA component extracted from the data can then be used for further analyses or simply plotted for visual inspection (Fig. 4).

**Figure 4:**
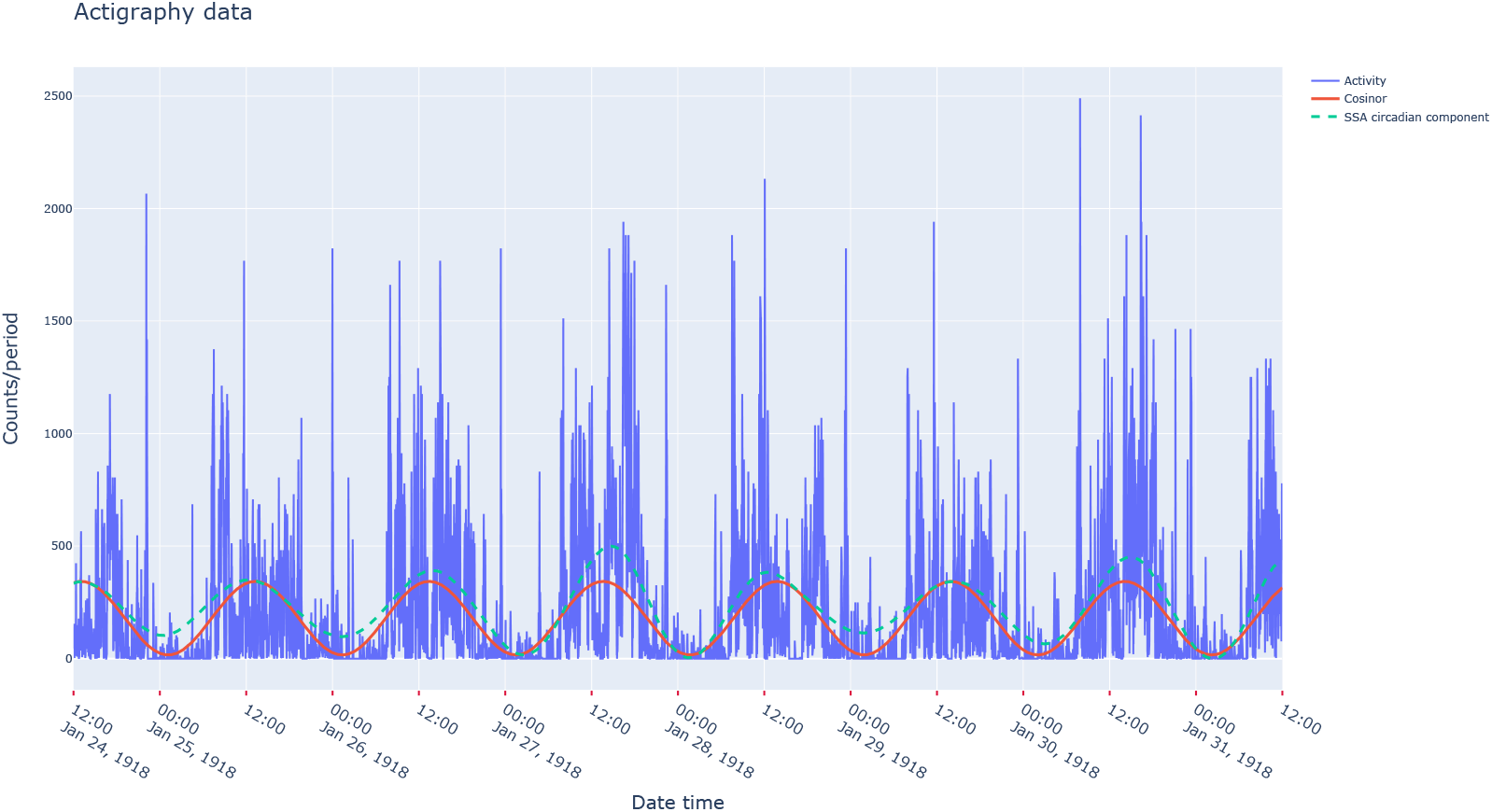
Vizualisation example.

More generally, complete informations on each function can accessed through the usual python “help” command or through the online documentation.

## 4. Impact

Even though actigraphy has been used in the field of sleep and chronobiology research for the past 40 years, there is, to our knowledge, no comprehensive open-source analysis package for actigraphy data that would allow users to read various data format, perform the necessary data cleaning as well as some more advanced data analysis within a single framework. This is all the more necessary as it would improve the reproducibility of research outcomes by limiting the proliferation of private analysis codes [35]. It would also allow users to perform more complex analyses and therefore make optimal use of actigraphy data that are often part of costly multi-modal data acquisition protocols. Such analysis package would also help to reduce error rates by alleviating the burden of manual data processing that hampers the processing of large-scale actigraphy datasets. The emergence of nation-wide biobanks, which would be crucial for understanding public health issues such as the impact of daylight time saving changes or chronic sleep deprivation, should be matched by the emergence of appropriate analysis tools. Besides, facilitating the access to such analysis tools for actigraphy data would benefit other fields of neuroscience. For example, there are evidence for a link between human brain structure and the locomotor activity, whether it is the total amount of activity [36, 37], the sleep fragmentation [38] or the integrity of the circadian rhythmicity [3, 39]. Human brain functions are also modulated by circadian and/or seasonal rhythmicity [40, 41]. Therefore, a precise assessment of rhythmicity, as allowed by actigraphy, is crucial for functional brain imaging and cognitive studies too. This is one of many examples that emphasize the benefit of extending the use of actigraphy outside the field of sleep and circadian research.

## 5. Conclusions

We present the *pyActigraphy* toolbox, an open-source python package for actigraphy data visualisation and analysis which offers functionalities to automatise data pre-processing, read large file batches and implement various metrics and techniques for the analysis of actigraphy data. By developing the *pyActigraphy* package, we not only hope to facilitate data analysis but also foster research using actimetry and drive a community effort to improve this open-source package and develop new variables and algorithms.

## Acknowledgements

The development of the *pyActigraphy* toolbox is part of the *CogNap* project that has received funding from the European Research Council (ERC) under the European Union’s Horizon 2020 research and innovation programme (Grant agreement No. 757763). This work was also supported by the Fonds de la Recherche Scientifique - FNRS under Grant n T.0220.20. CS is a research associate and MD is a FRIA grantee of the Fonds de la Recherche Scientifique FNRS, Belgium.

GH would like to warmly thank A. Lim and E. Winnebeck for having shared both insights and their code for the state transition probability and the LIDS analyses, respectively. GH would also like to thank A. Aubin and B. Leroy, who participated in the implementation of the *pyActigraphy* package during their summer internship at the GIGA-CRC *in vivo imaging* laboratory in 2018 and 2019, respectively.

## Current executable software version

Ancillary data table required for sub version of the executable software: (x.1, x.2 etc.) kindly replace examples in right column with the correct information about your executables, and leave the left column as it is.

**Table 2:**
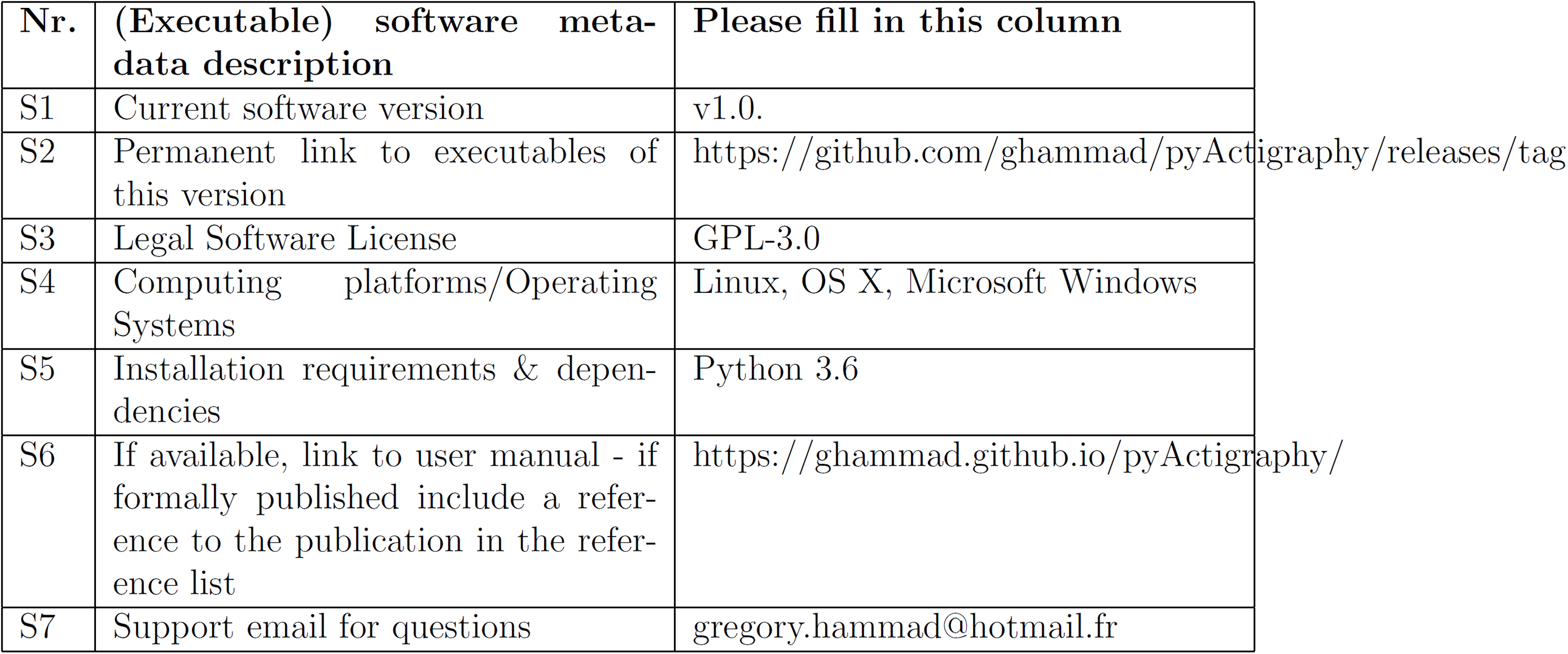
Software metadata (optional)

